# Safe and easy evaluation of tmRNA-SmpB-mediated *trans*-translation in ESKAPE pathogenic bacteria

**DOI:** 10.1101/2020.12.16.423090

**Authors:** Marion Thépaut, Rodrigo Campos Da Silva, Eva Renard, Frédérique Barloy-Hubler, Eric Ennifar, Daniel Boujard, Reynald Gillet

## Abstract

Bacteria cope with ribosome stalling thanks to *trans*-translation, a major quality control system of protein synthesis that is mediated by tmRNA, an hybrid RNA with properties of both a tRNA and an mRNA, and the small protein SmpB. Because *trans*-translation is absent in eukaryotes but necessary for bacterial fitness or survival, it is a promising target for the development of novel antibiotics. To facilitate screening of chemical libraries, various reliable *in vitro* and *in vivo* systems have been created for assessing *trans*-translational activity. However, none of these permits the safe and easy evaluation of *trans*-translation in pathogenic bacteria, which are obviously the ones we should be targeting. Based on green fluorescent protein (GFP) reassembly during active *trans*-translation, we have created a cell-free assay adapted to the rapid evaluation of *trans*-translation in ESKAPE bacteria, with 24 different possible combinations. It can be used for easy high-throughput screening of chemical compounds as well as for exploring the mechanism of *trans*-translation in these pathogens.

## Introduction

The World Health Organization (WHO) designated six ‘ESKAPE’ pathogens *(Enterococcus faecium, Staphylococcus aureus*, Klebsiella pneumoniae, *Acinetobacter baumannii, Pseudomonas aeruginosa*, and *Enterobacter spp*.) as critical targets for drug discovery (Rice et al., 2008; Tacconelli and Magrini, 2017). Indeed, these bacteria are the leading cause of nosocomial infections throughout the world, and most are multidrug-resistant isolates (Santajit and Indrawattana, 2016). The WHO recommendation is to focus specifically on the discovery and development of new antibiotics that are active against multidrug- and extensively drugresistant ESKAPE bacteria. However, the hazardous nature of these pathogens makes it highly challenging to develop high-throughput screening methods for identifying and evaluating any new antimicrobial agents for future clinical use. To aid in this, the molecular process to be targeted must first be identified, and ideally this process should be: i) conserved among all pathogenic ESKAPE bacteria; ii) indispensable to bacterial survival or at least its fitness; iii) sufficiently variable that different species can be distinguished from each other; iv) absent in eukaryotes; v) not targeted by current antibiotics; vi) unrelated to existing resistance mechanisms; and finally vii) reproducible in non-hazardous *in vitro* experiments.

In fact, *trans*-translation appears to be the perfect candidate. This mechanism is the primary bacterial rescue system, allowing for the release of ribosomes stalled on faulty mRNAs that lack stop codons as well as the elimination of these mRNAs and mistranslated peptides. The *trans*-translation process is performed by hybrid transfer-messenger RNA (tmRNA) and its protein partner SmpB (Giudice *et al.*, 2014). Briefly, the tmRNA-SmpB complex recognizes the stalled ribosome and associates with it. In a finely orchestrated ballet, translation then resumes on tmRNA’s internal mRNA-like domain (MLD), which encodes a specific sequence that is recognized by proteases. This process permits the stalled ribosomes to be recycled, the degradation of the incomplete peptide after its release, and elimination of the problematic nonstop mRNA. Remarkably, genes coding for tmRNA and SmpB have been found in nearly all bacterial genomes, yet not in eukaryotes, with the exception of a very few rare organelles (Hudson and Williams, 2015). Despite high sequence conservation at both the 5’- and 3’-ends of tmRNA genes, the internal sequences of tmRNA are considerably divergent among different species (Supp. Fig. 1), and this property makes tmRNA a good tool for species identification (Schonhuber *et al.*, 2001). In the same way, despite global structural conservation, variations in *smpB* sequences are also considerably divergent among different species (Supp. Fig. 1).

While resolving stalled ribosomal complexes is undoubtedly a matter of life or death (Keiler and Feaga, 2014), *trans*-translation itself is not always indispensable to bacterial survival. This irregularity was the subject of a long debate until the discovery of backup systems, mechanisms which take over if *trans*-translation is deficient or overwhelmed. However, even when they are present, these systems are not enough to ensure a steady and prolonged fitness to the cell, as impaired *trans*-translation is known to result in various phenotypes varying from mild (such as loss of tolerance to multiple antibiotics and stresses) to severe (including lethality or loss of virulence) (Li *et al.*, 2013; Keiler and Feaga, 2014). To date, *trans*-translation has not been yet exploited for clinical use. In the search for inhibitors specific to the process, initial assays led to the discovery of 1,3,4 oxadiazole molecules (Ramadoss *et al.*, 2013), but their activity *in vivo* is still in question (Macé *et al.*, 2017; Brunel *et al.*, 2018). It has been suggested that *trans*-translation is inhibited by pyrazinamide (PZA), a first-line anti-tuberculosis drug (Shi *et al.*, 2011), but it was finally recently shown the action of PZA is entirely independent of *trans*-translation in *M. tuberculosis* (Dillon *et al.*, 2017).

Because of its biological properties, transposition of *trans*-translation into a non-hazardous system that could allow for rapid and easy evaluation of its activity would greatly help in the search for new antibiotics which target this system. While there are routine methods for screening the antimicrobial activity of compounds from chemical libraries, a combination of this primary screening with the specification of a molecular target is much harder to implement (Osterman *et al.*, 2016). An ideal method would allow not just the identification of the targeted cellular process, but also its level of specificity toward a bacterial genus or species. Furthermore, an easy quantitative and rapid analysis of the process should be possible even in small volumes. Reporter assays are the best candidates for efficient initial high-throughput screening (HTS) methods, as they can be quick and automated, as well as quite useful for screening unpurified mixtures of natural extracts (Osterman *et al.*, 2016). Accordingly, we recently used a commercial reconstituted *in vitro* translation system (PURExpress) to create a reliable *in vitro* reporter system that detects the *E. coli trans*-translation activity (Guyomar *et al.*, 2020). This assay, based on reassembling an active “superfolder” Green Fluorescent Protein (sfGFP) after tmRNA tagging (Fig. 1), was designed and validated for the specific *in vitro* quantification of *trans*-translation in ESKAPE pathogenic bacteria, and we report on that here.

**Figure 1.**
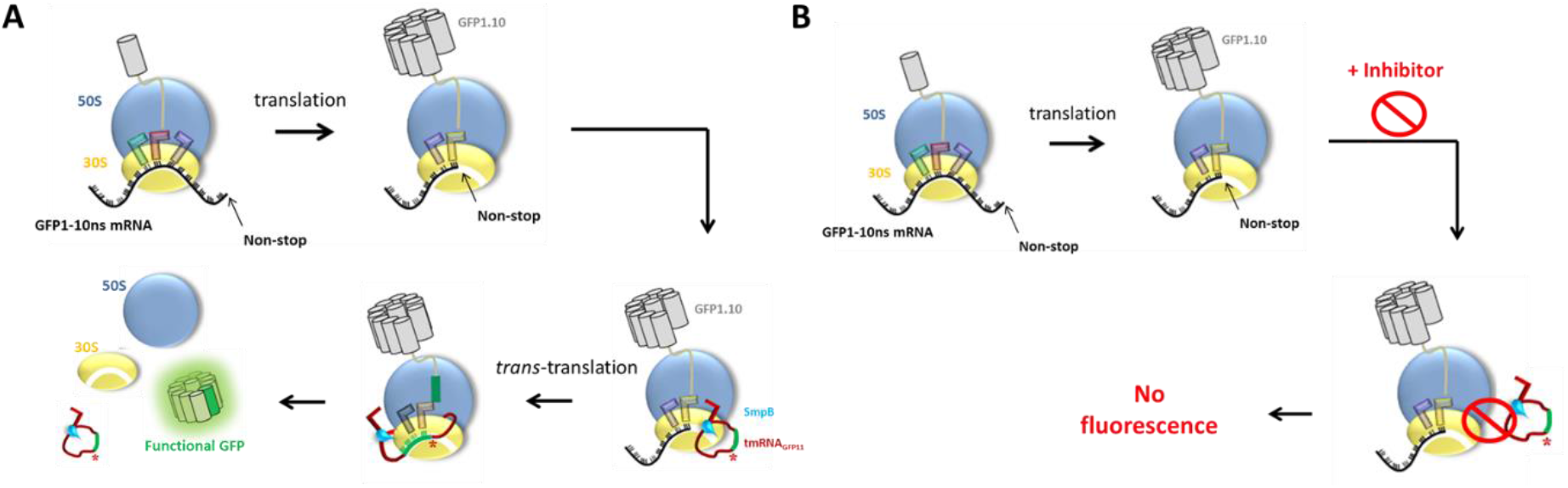
**A. *Trans*-translation of sfGFP1-10 mRNA lacking a stop codon.** The tmRNA_GFP11_-SmpB complex binds to the stalled ribosome, and translation resumes thanks to the tmRNAGFP 11 mRNA-like domain (MLD). The MLD encodes the missing eleventh domain of the sfGFP, and the complete sfGFP is released and becomes fluorescent. **B. Impairment of the process in the presence of *trans*-translation inhibitors**. The ribosomes stay stalled on the problematic mRNA and fluorescence is impaired.

## Results

### Distribution of ArfA and ArfB in ESKAPE bacterial genomes

If some bacteria can live without *trans-translation*, this is only because they have at least one of the two alternative release factors, ArfA and ArfB (Himeno *et al.*, 2015). Indeed, the ArfA or ArfB backup systems are essential if *trans*-translation is aborted. Depending on backup system status, therefore, the effects of specific inhibitors of *trans*-translation will vary - from increasing the activity of currently used antibiotics to outright cell death. It was therefore important for us to begin by pinpointing the phylogenetic distribution of ArfA and ArfB in ESKAPE pathogens. To do this, we investigated the sequences of those rescue factors using a combination of *in silico* methods including keyword searches, similarity detection, protein domain prediction, ortholog clustering, and synteny analysis. This pipeline was applied to the complete genomic sequences of 1,670 species: 148 *E. faecium*, 459 *S. aureus*, 465 *K. pneumoniae*, 188 *A. baumannii*, 259 *P. aeruginosa*, and 151 *Enterobacter* spp. Interestingly, among these ESKAPE pathogens, neither of the two back-up systems were found in *A. baumannii* or the two Gram-positive bacteria *E. faecium* and *S. aureus* (Table 1). While we cannot categorically state that no backup systems exist in these bacteria - see for instance the recent reports on ArfT in *Francisella tularensis* and BrfA in *Bacillus subtilis* (Goralski *et al.*, 2018; Shimokawa-Chiba *et al.*, 2019) - we can however suppose that their viability depends on *trans*-translation impairment. On the other hand, we found genes encoding ArfA and/or ArfB in most if not all of the *K. pneumonia, P. aeruginosa*, and *Enterobacter* spp. studied. The impairment of *trans*-translation in these bacteria is probably less severe, therefore, even if it still detrimental to bacterial fitness.

**Table 1.** Distribution of ArfA and ArfB in ESKAPE bacteria

| ESKAPE pathogen | # of screened genomes | *arfA* | *arfB* |
| --- | --- | --- | --- |
| *Enterococcus faecium* | 148 | 0 | 0 |
| *Staphylococcus aureus* | 459 | 0 | 0 |
| *Klebsiella pneumoniae* | 465 | 464 + 1 $\Delta$ | 459 + 6 $\Delta$ |
| *Acinetobacter baumannii* | 188 | 0 | 0 |
| *Pseudomonas aeruginosa* | 259 | 259 | 259 |
| *Enterobacter* spp. | 151 | 151 | 150 + 3 $\Delta$ |

$\Delta$ pseudogenes with frameshift or « in-frame » stop codon.

### ESKAPE tmRNA and SmpB production

To allow for independent monitoring of *trans*-translation in these six ESKAPE pathogens, we engineered their tmRNAs by replacing their internal MLD with a sequence of 16 amino acids that encodes GFP’s eleventh domain (Supp. Table 1). To conserve the tmRNA H5 helix that is instrumental during *trans*-translation, we also engineered compensatory mutations (Guyomar *et al.*, 2020, and Fig. 2A, B). Unlike those of the other bacteria, the natural tmRNA 3’-ends in *E. faecium* and *S. aureus* are not CCA but UUG and UAU, respectively, so these were replaced by CCA to ensure correct aminoacylation by *E. coli* AlaRS (Barends *et al.*, 2000), and these variants were named tmRNA_GFP11_.

**Figure 2.**
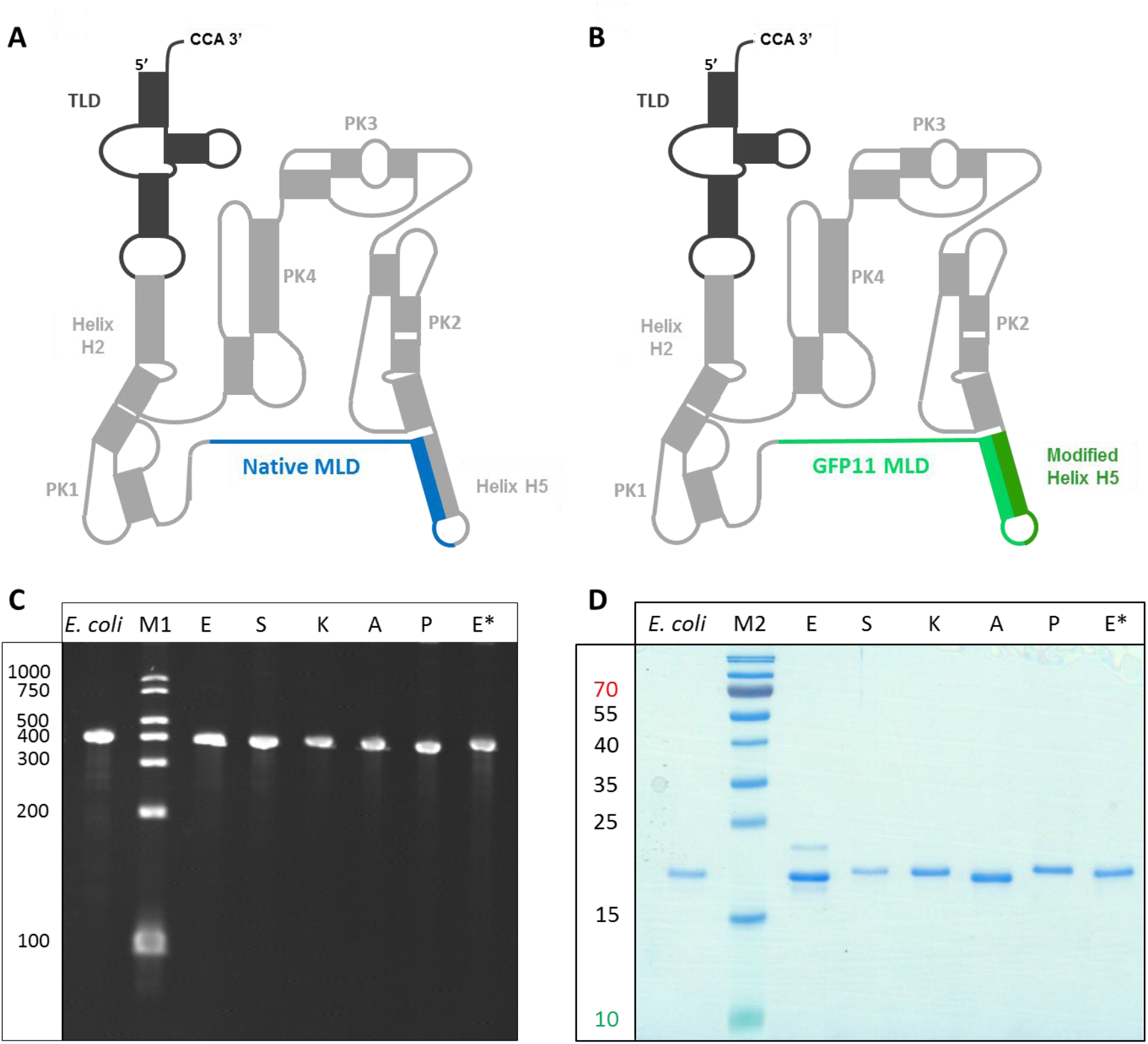
Schematic illustrations of tmRNA secondary structures as well as production patterns of tmRNA and SmpB variants. (A) Wild-type tmRNA, with a black transfer-like domain (TLD) and blue messenger-like domain (MLD). (B) Mutated tmRNAoFPu has an engineered MLD (light green) that encodes the eleventh GFP domain beta-strand. Compensatory mutations (dark green) maintain the base-pairing interactions of the H5 helix, and the 3’-ends for all species are CCA (Adapted from Guyomar *et al.*, 2020.) (C) Visualization of ESKAPE tmRNA_GFP11_ variants (5 pmol) on 8% urea-PAGE, with *E. coli* tmRNA_GFP11_ used as a control. (D) As C, but showing 50 pmol *E. coli* and ESKAPE SmpB on 15% SDS-PAGE. Key: M1, RNA Century™-Plus Markers (ThermoFisher Scientific), M2, PageRuler™ Prestained Protein Ladder, 10 to 180 kDa (ThermoFisher Scientific), E, *Enterococcus faecium;* S, *Staphylococcus aureus;* K, *Klebsiella pneumoniae;* A, *Acinetobacter baumannii;* P, *Pseudomonas aeruginosa*; E*, *Enterobacter cloacae*.

Urea-PAGE analysis indicated that the six tmRNA variants were successfully produced at the expected size, without any noticeable degradation or unexpected bands (Fig. 2C). We started with 10 μg plasmidic DNA, and the final yields were about 4 nmol of transcribed RNAs for each reaction. The six corresponding SmpB proteins were cloned and produced *in vivo* in *E. coli* (see Experimental Procedures). After purification, acrylamide analysis confirmed the correct size of each protein (Fig. 2D). The final yields for each ESKAPE SmpB were about half the amount of the *E. coli* SmpB produced.

### ESKAPE trans-translation reactions

In order to obtain non-productive translation complexes (NTCs) to be targeted by *trans*-translation, we used a reconstituted cell-free protein synthesis (NEB PURExpress) system from *E. coli* (Shimizu, *et al.*, 2001, Shimizu and Ueda, 2002). By adding a non-stop DNA template, we were able to accumulate stalled ribosomes with the ten first domains of sfGFP stuck in the ribosome exit tunnel (Fig. 1A). When tmRNA_GFP11_ and *E. coli* SmpB are added, the ribosomes are freed and the intensity of the fluorescent signal increases over time while the complete sfGFP protein is produced. A plateau is reached at ~4 hours of incubation, and the fluorescence remains stable for at least 710 minutes (Fig. 3A, black curve).

**Figure 3:**
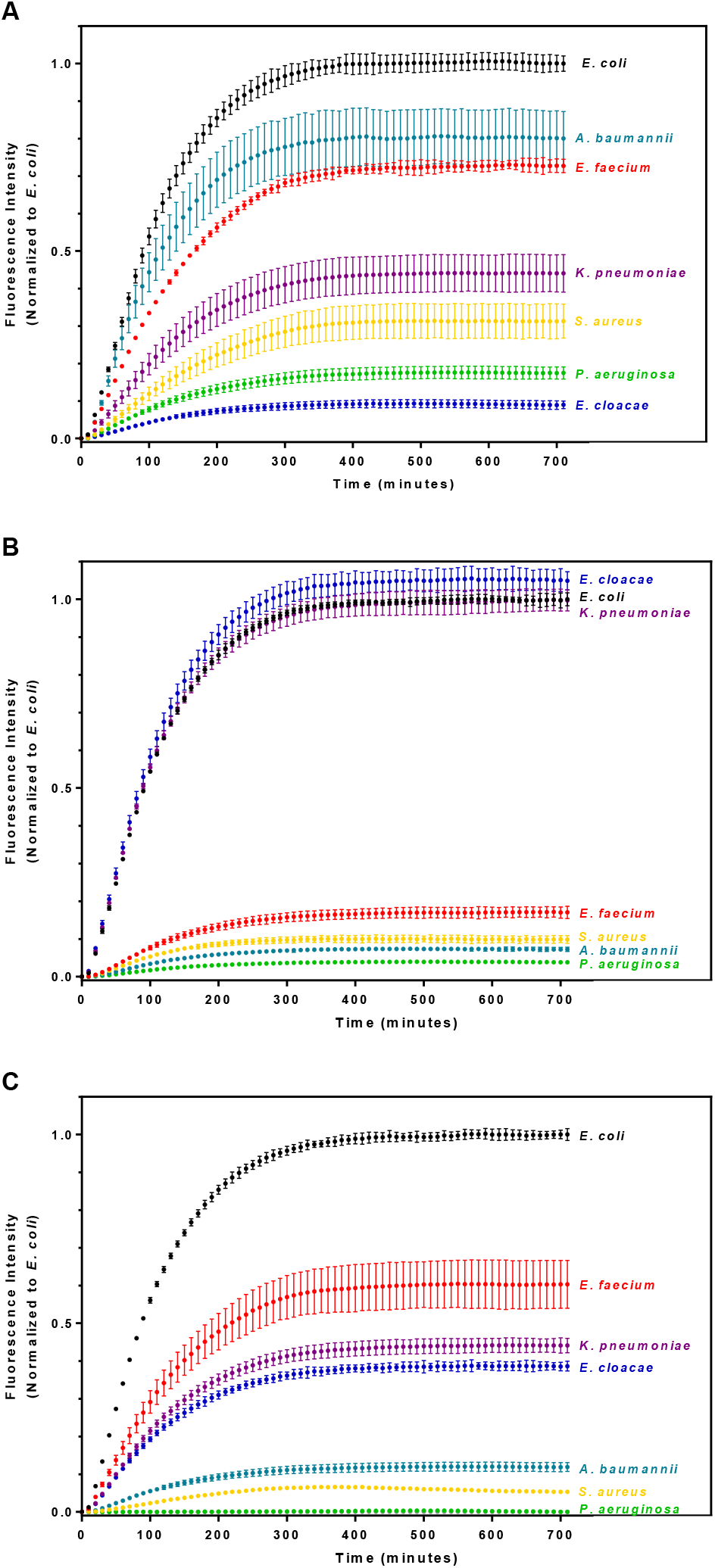
*Trans*-translation kinetics over time using *Escherichia coli* ribosomes. Fluorescence increases are directly linked to *trans*-translation activity. **A)** *Trans*-translation assays were done on *E. coli* tmRNA_GFP11_ using the SmpBs from each ESKAPE pathogen, with the *E. coli* SmpB as a control. **B)** *Trans*-translation assays keeping the *E. coli* SmpB but using the tmRNA_GFP11_ variants of each ESKAPE pathogen, with an *E. coli* tmRNA_GFP11_ as the control. **C)** Both SmpB and tmRNA_GFP11_ are from each ESKAPE pathogen, with the *E. coli* SmpB-tmRNA_GFP11_ as a control. The results are shown as means ± standard deviation and normalized to the *E. coli* conditions.

In a first set of heterologous experiments, we kept the *E. coli* tmRNA_GFP11_, but replaced its SmpB by one from an ESKAPE pathogen. A fluorescent signal was still recovered with each one of the hetero-complexes, albeit at different levels (Fig. 3A). The *E. cloacae, S. aureus*, and *P aeruginosa* SmpBs displayed the lowest signals, less than 30% of the *E. coli* control, while the *K. pneumoniae* SmpB signal was about half the control, and *E. faecium* and *A. baumannii* at 80%. This demonstrates that all of the ESKAPE SmpBs are functional and sufficiently conserved to be interchangeable in the presence of *E. coli* tmRNA. While it confirms that SmpB is highly conserved (Supp. Fig. 1), it also supports the use of this simple system for screening molecules that target SmpB but not tmRNA. Indeed, since SmpB is essential for tmRNA’s peptide-tagging activity (Karzai *et al.*, 1999), disrupting SmpB is one of the most promising ways to impair *trans*-translation. In fact, aptamers that inhibit SmpB functioning were recently shown to trigger strong growth defects in *Aeromonas veronii* C4 (Liu *et al.*, 2016).

We then performed the experiments the other way around, using the *E. coli* SmpB but the tmRNAs from the ESKAPE pathogens. Contrary to the previous experiments, only the heteroduplexes combining *E. coli* SmpB and the tmRNAs from *K. pneumoniae* and *E. cloacae* gave out strong signals, about the same levels as those recovered in the *E. coli* tmRNA control (Fig. 3B). This is not a surprise since, like *E. coli*, both *K. pneumoniae* and *E. cloacae* are *Enterobacteriaceae* with very similar tmRNA sequences (≥ 95% identity with *E. coli*, see Supp. Fig. 1). The four other bacterial species all produced signals, but at lower levels (about 5 to 20% of the reference).

We continued by performing homologous experiments, using SmpB and tmRNA_GFP11_ from the same ESKAPE pathogen, but this time with *E. coli* ribosomes (Fig. 3C). Five of the six complexes yielded positive results. Three of these were at high levels (~50% compared to the *E. coli* reference): *K. pneumonia*; *E. cloacae* and *E. faecium*. The other two were at lower levels (about 5-10% of the reference): *A. baumannii*, another Gammaproteobacteria that is relatively close to *Enterobacteriaceae;* and the Gram-positive *S. aureus.* The Gammaproteobacteria *P. aeruginosa* did not work at all.

In a final set of experiments, we used homologous tmRNA-SmpB complexes in the presence of their corresponding ESKAPE ribosomes. The use of the PURExpress Δ Ribosome kit allowed us to substitute commercial *E. coli* ribosomes with ESKAPE variants we had prepared in-house. We first confirmed the effectiveness of translation using these ribosomes by synthesizing full-size sfGFP. For all ribosomes used, a fluorescent signal was recovered, indicating that the ESKAPE ribosomes translate well even if at lower levels (Fig. 4A). The *P. aeruginosa* and *E. cloacae* Enterobacteriaceae ribosomes gave the strongest signals, around 45 and 35% respectively as compared to that of *E. coli.* All of the other signals were below 20%, even dropping under 10% in the case of *S. aureus.*

**Figure 4.**
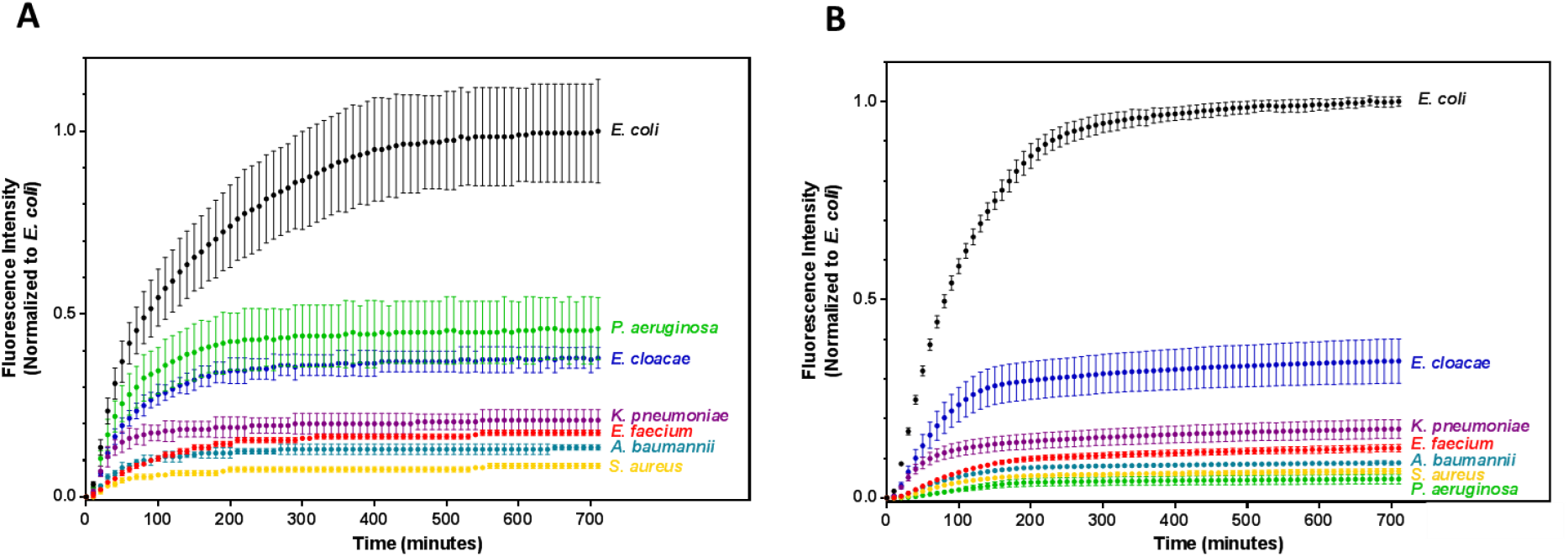
Translation and *trans*-translation kinetics over time. **A) Translation kinetics over time**: the increase in fluorescence is directly linked to translation. **B) *Trans*-translation kinetics over time using ESKAPE ribosomes**. All results are shown as means ± standard deviation and normalized to *E. coli.*

Despite these rather poor translation rates, fluorescence was easily detected, so we also performed *trans*-translation experiments using ESKAPE ribosomes (Fig. 4B). The goal was to improve the levels of the *trans*-translation signals previously recovered, but more importantly to obtain a positive result for *P. aeruginosa.* The results were finally conclusive for that bacteria, which gave a fluorescent signal of ~10% compared to the control. This positive result is certainly due to the quite efficient translation obtained with these ribosomes (Figure 4A). On the other hand, the *trans*-translation levels of the other bacteria did not improve, and were even lower in *S. aureus*. This is surely due to the fact that the PURExpress system is based on only *E. coli* translation factors, and their low count limits their handling of canonical translation (see Fig. 4A) or specific tmRNAs *(e.g.* tmRNA aminoacylation by *E. coli* AlaRS or tmRNA-SmpB transport by *E. coli* EF-Tu-GTP).

## Discussion

Here we describe the use of GFP as a reporter for safe measurement of the *trans*-translation activity of the six ESKAPE systems in a cell-free protein synthesis system. The various combinations we evaluated (four for each ESKAPE pathogen) have yielded different interesting strategies for the disruption of *trans*-translation (Fig. 5).

**Figure 5:**
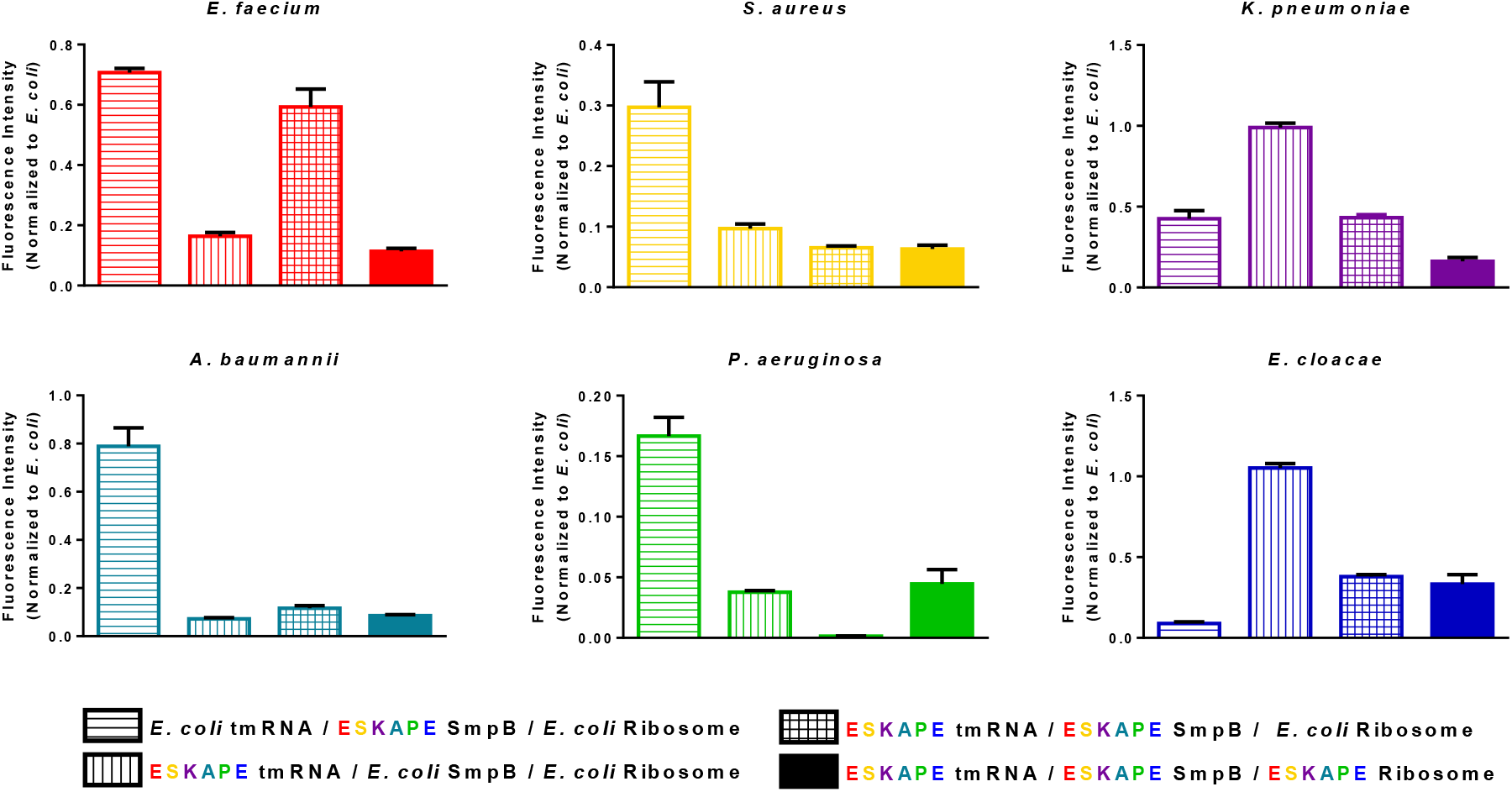
Quantification of *in vitro trans*-translation. Normalized fluorescence obtained in heterologous and homologous systems are shown at 310 minutes of incubation and reassembled by species. The results were normalized to the *E. coli* conditions and are shown as means ± standard deviations.

The molecules being investigated for the development of new anti-*trans*-translation antibiotics will have different ways of interfering with tmRNA-SmpB binding to stalled ribosomes. They could disrupt tmRNA-SmpB interactions, or they could prevent interactions between the complex and the ribosome, such as by blocking the entrance of SmpB entirely or by preventing the passage of the complex through the bridges which have to be open during the process. Therefore, it is of great interest to have the ability to evaluate the targeting of the three main actors (tmRNA, SmpB, and the ribosome) independently as well as in each ESKAPE system. Of the 24 combinations we tested, 23 exhibited a signal strong enough for evaluating the possible activity of inhibitors. The only one that did not was the *P. aeruginosa* tmRNA-SmpB complex when used with *E. coli* ribosomes. We first suspected that the tmRNA H5 helix, inspired from the *E. coli* helix (Fig. 1B), might somehow have altered its activity. Therefore, to avoid any possible effects of the helical rearrangement, we constructed and tested new tmRNA_GFP11_ variants for *P. aeruginosa* but also *E. coli, S. aureus* and *E. faecium.* These tmRNA_GFP11_V2 constructs all have the full sequence that encodes the eleventh domain of GFP upstream from the natural H5 helix (Supp. Fig. 2). However, these variants did not have improved fluorescence, and *P. aeruginosa* still did not emit signals. We can thus exclude the idea that the different structural features between the *P. aeruginosa* and *E. coli* ribosomes (Halfon *et al.*, 2019) are important enough to prevent the correct process from occurring.

To permit the high-throughput screening of chemical compounds in multi-well microplates it was important to lower average screening costs of the current assay. To enable this, we decreased the reaction scale of the assays by reducing the final reaction volumes down to a microliter scale. Proof-of-concept experiments were performed with *E. coli* in final volumes of 1 μl using a MANTIS liquid-handler instrument (Formulatrix), and the resulting signals were strong enough to allow for the easy detection of *trans*-translational activity (not shown). Indeed, the objective of this study was to create a non-hazardous *in vitro* screening system for evaluating *trans*-translation in ESKAPE pathogens, and to miniaturize it for HTS applications, and the assays we performed were convincing. The system will clearly be very effective for benchmarking the effects of new antibiotic compounds that target *trans*-translation in highly pathogenic bacteria, as well as aiding us to better understand the *trans*-translation process in these bacteria. Its flexibility in the choice of target bacterial species and the possibility to for varying the combinations of tmRNA, SmpB, and ribosomes are advantageous, making the identification of new specific antimicrobial inhibitors easier. Ongoing experiments in our laboratory are using this to screen large chemical and natural product libraries for drug discovery.

## Experimental Procedures

### *In silico* analysis

Complete genomes were retrieved from the NCBI database (March 2020). Chromosomes and plasmids (when present) were studied separately. GenBank files were first searched based on their textual annotation entries, using the keywords ‘ArfA,’ ‘yhdL,’ and ‘alternative ribosome-rescue factor’ (for ArfA), or ‘ArfB,’ ‘yaeJ,’ ‘ribosome-associated protein,’ and ‘peptidyl-tRNA hydrolase’ (for ArfB). Missing loci were checked using BlastN, BlastP, and tBlastN similarity-detection strategies (Altschul *et al.*, 1990) as well as comparative genomics, with synteny analysis done using progressiveMauve (Darling *et al.*, 2010). All retrieved loci were compared using the Reciprocal Best Hits method, and InterProScan (Jones *et al.*, 2014) was used on the corresponding proteins to check for the presence of the IPR005589/ PF03889 (ArfA) and IPR000352/PF00472 (ArfB) domains. Frameshifted loci were indicated as annotated in the GenBank files. Finally, the presence and absence of K09890 (ArfA) and K15034 (ArfB) were checked in the KEGG ORTHOLOGY database (Kanehisa *et al.*, 2016).

### Plasmid construction and preparation

For each ESKAPE tmRNA, the internal open reading frame was replaced by the eleventh domain of the superfolder GFP (sfGFP) preceded by the first conserved alanine of native tmRNA. In order to preserve the H5 helix, compensatory mutations were added (Fig. 2B).

Additionally, the sequences were designed to carry a T7 promotor sequence in the 5’-end in order to realize transcription *in vitro.* Note that the tmRNA 3’-end natural sequences from *E. faecium* (UUG) and *S. aureus* (UAU) were replaced by CCA so that the *E. coli* AlaRS could correctly aminoacylate them.

We also produced tmRNA_GFP11_V2 variants for *E. coli, P.aeruginosa, S.aureus* and *E. faecium* species. This tmRNA_GFP11_ series carries the full sequence encoding the eleventh domain of GFP upstream from the *E. coli* H5 helix (Supp. Fig. 2). In order to obtain mature tmRNA_GFP11_ by *in vitro* transcription, the tmRNA_GFP11_ and tmRNA_GFP11_V2 ESKAPE sequences were synthesized and cloned into the pUC19 vector between the HindIII and BamHI restriction sites (Supp. Table 1). For each ESKAPE SmpB, GenScript synthesized the sequences with codon optimization for *E. coli*, cloning them into the pET22b(+) vector between the NdeI and XhoI restriction sites to add a 6His histidine tag (Supp. Table 2). The generated plasmids, pUC19ESKAPEtmRNA_GFP11_ and pET22b+ESKAPESmpB (Supp. Table 5), were amplified in *E. coli* NM522 cells then extracted using a NucleoBond Xtra Midi kit (Macherey-Nagel). Quantification was performed using a SimpliNano spectrophotometer (Biochrom).

### SmpB purification

The bacterial cultures and SmpB purification were all done as previously described (Guyomar *et al.*, 2020). His-tagged *E. coli* and ESKAPE SmpB proteins (Supp. Table 2) were expressed from the pF1275 and the pET22b+ESKAPE SmpB vectors under the control of a T7 promoter in BL21(DE3)*ΔssrA* cells (Cougot *et al.*, 2014). Briefly, BL21(DE3)Δ*ssrA* cells were grown in lysogeny broth (LB) at 30 °C supplemented with ampicillin (100 μg/ml) and kanamycin (50 μg/ml). Protein expression was induced in the exponential phase *(OD_600_* = 0.6) with 0.1 mM isopropyl-β-D-1-thiogalactopyranoside (IPTG) overnight at 16 °C. Cells were harvested and washed, then resuspended in lysis buffer (50 mM HEPES-KOH, 200 mM KCl, 20 mM imidazole, and 1 mM DTT pH 7.5). Cell lysis was performed using a French press, and the lysate was centrifuged at 15,000 rpm for 45 minutes at 4 °C in a Beckman J2-MC with a JA-17 rotor. The supernatant was then filtrated (0.2 μm) and injected onto a Ni-NTA sepharose column (HisTrap FF, GE Healthcare) previously equilibrated with the lysing buffer. The column was washed with 100 ml lysis buffer and 50 ml washing buffer (50 mM HEPES-KOH, 200 mM KCl, 1 M NH4Cl, imidazole 20 mM, and 1 mM DTT pH 7.5) before elution with 500 mM imidazole. Finally, a 10kDa Amicon Ultra centrifugal filter (Merck Millipore) was used to concentrate the fractions containing pure SmpB, changing the buffer to a concentration buffer (50 mM HEPES-KOH, 100 mM KCl, 10% glycerol, and 1 mM DTT pH 7.5). In order to visualize SmpB, 50 pmol of denatured proteins was analyzed on 15% SDS-PAGE gels. Proteins were detected using InstantBlue protein stain (Expedeon) according to the supplier’s instructions.

### tmRNA_GFP11_ production

*E. coli* and ESKAPE tmRNA_GFP11_ were produced as previously described (Guyomar *et al.*, 2020). Each ESKAPE tmRNA_GFP11_ was transcribed *in vitro* from the pUC19ESKAPEtmRNA_GFP11_ plasmids. To generate the 3’ end needed for aminoacylation by AlaRS, the plasmid (10 μg) was completely digested by NEB BsmBI or EarI restriction enzymes (Supp. Table 5). After phenol/chloroform extraction, the purified digested plasmid was precipitated, and the resulting pellets resuspended in 40 μl nuclease-free water. A MEGAscript T7 transcription kit (Thermo Fisher Scientific) was used to produce each ESKAPE tmRNA_GFP11_ before its purification using the corresponding MEGAclear kit. Denatured tmRNA_GFP11_ was checked by electrophoresis on 8% Urea-PAGE gels, stained with ethidium bromide, and visualized under ultraviolet light.

### DNA templates and oligonucleotide production

For *trans*-translation assays, the nonstop GFP1-10 sequence was produced by PCR using primers #1 and 2 and Q5 High-Fidelity DNA Polymerase (NEB) with pETGFP 1-10 vector as a template (Cabantous *et al.*, 2006; Supp. Table 3 and 4). For translation assays, primers #1 and #3 from the same template were used to amplify sfalaGFP, the superfolder GFP having an additional conserved alanine between the sfGFP1-10 and sfGFP11 domains (Supp. Table 3 and 4). The resulting PCR products were purified using a QIAquick PCR Purification Kit (Qiagen) and checked by agarose electrophoresis. Both PCR products have a T7 promoter upstream from their coding sequences. Antisense oligonucleotide “A” was supplied by Eurofins (Supp. Table 3).

### ESKAPE ribosome purification

Ribosomes were purified from *Acinetobacter baumannii* (clinical isolate); *Staphylococcus aureus* (clinical isolate); *Pseudomonas aeruginosa* (ATCC 27853); *Enterobacter cloacae* (clinical isolate); *Klebsiella pneumoniae* (clinical isolate); and *Enterococcus faecium* (HM1070). From an overnight starter culture, 6-9 L of LB medium were inoculated to reach an OD_600_ of 0.05, then stirred at 150 rpm at 37 °C. Bacterial growth was stopped when the OD_600_ reached 0.8 to 1.0. The cells were then centrifuged at 4,000 rpm for 20 minutes at 4 °C. Pellets (around 2 g/L of culture) were washed in a lysis buffer (20 mM Tris-HCl pH 7.5, 20 mM MgCl2, 200 mM NH4Cl, 0.1 mM EDTA, and 6 mM β-mercaptoethanol), centrifuged at 4,000 rpm for 15 minutes at 4 °C, and kept overnight at −80 °C. Pellets were then suspended in a Potter homogenizer in another lysis buffer complemented with 1 mM CaCl_2_. Cells were lysed in a French press at 1.0 kbar. To remove cellular debris, the lysates were centrifuged using a type 50.2 Ti rotor at 18,200 rpm for 30 minutes at 4 °C. The superficial pellet layer was then discarded, and the pellet resuspended in lysis buffer. Ribosomes were isolated by centrifuging lysates on a 30% sucrose cushion at 31,500 rpm for 19 hours at 4 °C. The superficial layer of pellets was again discarded, leaving only the transparent pellets which were then resuspended in conservation buffer (20 mM Tris-HCl pH 7.5, 10 mM MgCl2, 50 mM NH4Cl, 0.1 mM EDTA, and 6 mM β-mercaptoethanol). Any remaining contaminants were removed by a final centrifugation at 18,200 rpm for 1 hour at 4 °C. Ribosomes were concentrated using a Centricon (Merck Millipore) with a cut-off of 100K, flash-frozen in nitrogen, and conserved at −80 °C.

### *Trans*-translation assays

*In vitro trans*-translation assays were performed using the PURExpress *In Vitro* Protein Synthesis and Δ Ribosome kits (New England Biolabs). For *trans*-translation assays, PURExpress was supplemented by 62.5 ng purified PCR product encoding for nonstop sfGFP1-10, 12.5 pmol tmRNA_GFP11_, 25 pmol SmpB, and 50 pmol antisense A. Where necessary (Δ Ribosome), 6.725 pmol ribosomes were also added. These reactions were performed in a final reaction volume of 10 μL, with PURExpress diluted by a final factor of 1.6 with Buffer III (HEPES-KOH 5mM pH7.5, MgOAc 9mM, NH4Cl 10mM, KCl 50mM, and DTT 1mM). A Step One Plus PCR system (Applied Biosystems) was used for incubation at 37 °C as well as for fluorescence measurements over 710 minutes.

### Translation assays

*In vitro* translation assays were performed using a PURExpress Δ Ribosome kit. To produce the sfalaGFP, the PURExpress Δ Ribosome was diluted to a final factor of 1.6 with Buffer III, to which was added 62.5 ng purified PCR product and 6.725 pmol of the appropriate ribosomes. The translation reactions were incubated at 37 °C, and fluorescence was measured over 710 minutes using a Step One Plus.

## Acknowledgments

The authors are particularly grateful to Axel Innis and Anne-Xander van der Stel for sharing their Mantis experiment expertise and to Charlotte Guyomar for insightful comments on the manuscript. We thank Sylvie Georgeault and Fanny Demay for their help in purifying *E. faecium* ribosomes.

## Funding and additional information

This work was supported by the Agence Nationale pour la Recherche as part of the EU’s Joint Programming Initiative on Antimicrobial Resistance project (JPIAMR) project “Ribotarget - Development of novel ribosome-targeting antibiotics” as well as by SATT Ouest-Valorisation (DV 2552 and DV 3506).

## Author contributions

M.T. cloned and purified the SmpB and tmRNA variants and performed and analyzed the *trans*-translation *in vitro* assays. R.C.D.S. performed *trans*-translation *in vitro* assays. E.V. purified ribosomes from *A. baumannii, E. cloacae, K. pneumoniae, P. aeruginosa*, and *S. aureus.* F.B.H. performed *in silico* analysis of ArfA and ArfB in bacterial genomes. R.G. designed the study. E.E., D.B., and R.G. supervised the project. M.T. and R.G. wrote the manuscript, and all authors approved its final version.

## Supporting Information

### A - Phylogenetic comparison of tmRNA sequences

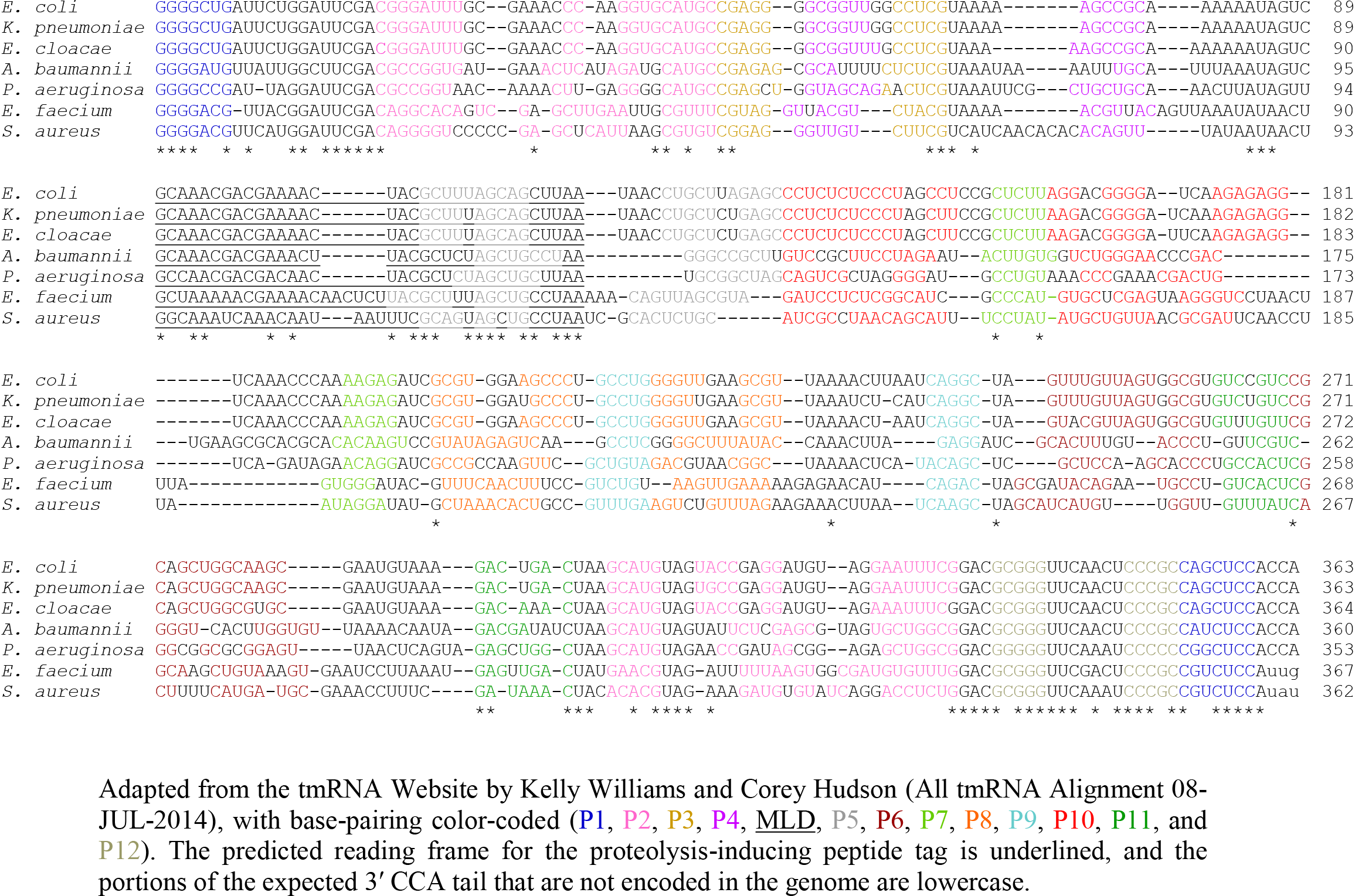

### A - Phylogenetic comparison of SmpB amino-acid sequences

**Supplementary Figure 1.**
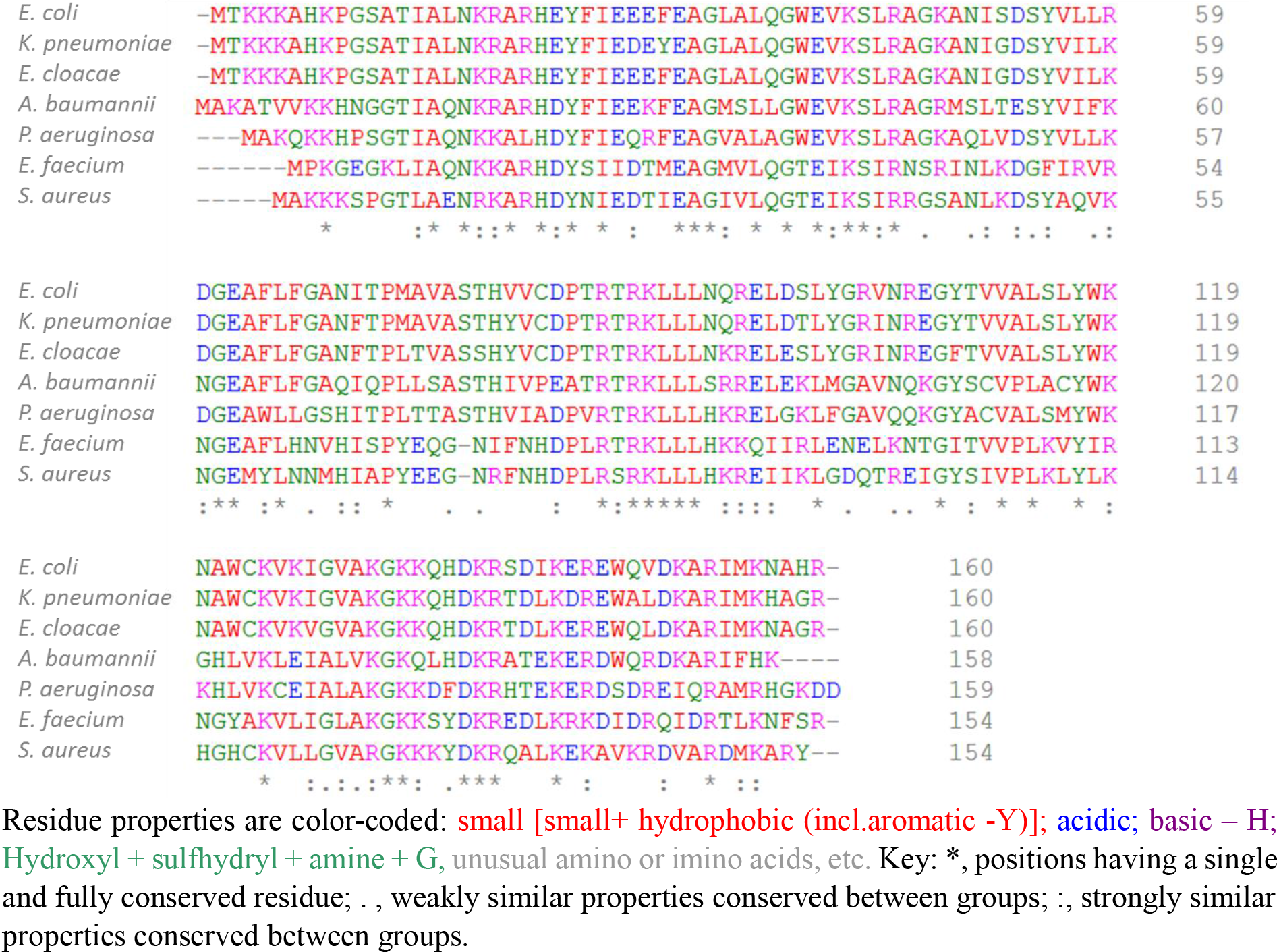
Phylogenetic comparison of SmpBs and tmRNAs from ESKAPE pathogens and *E. coli*. A) Shown here are the SmpB amino acid sequences (Clustal Omega) and B) the RNA sequences of tmRNAs from ESKAPE and *E. coli* bacteria.

**Supplementary Figure 2.**
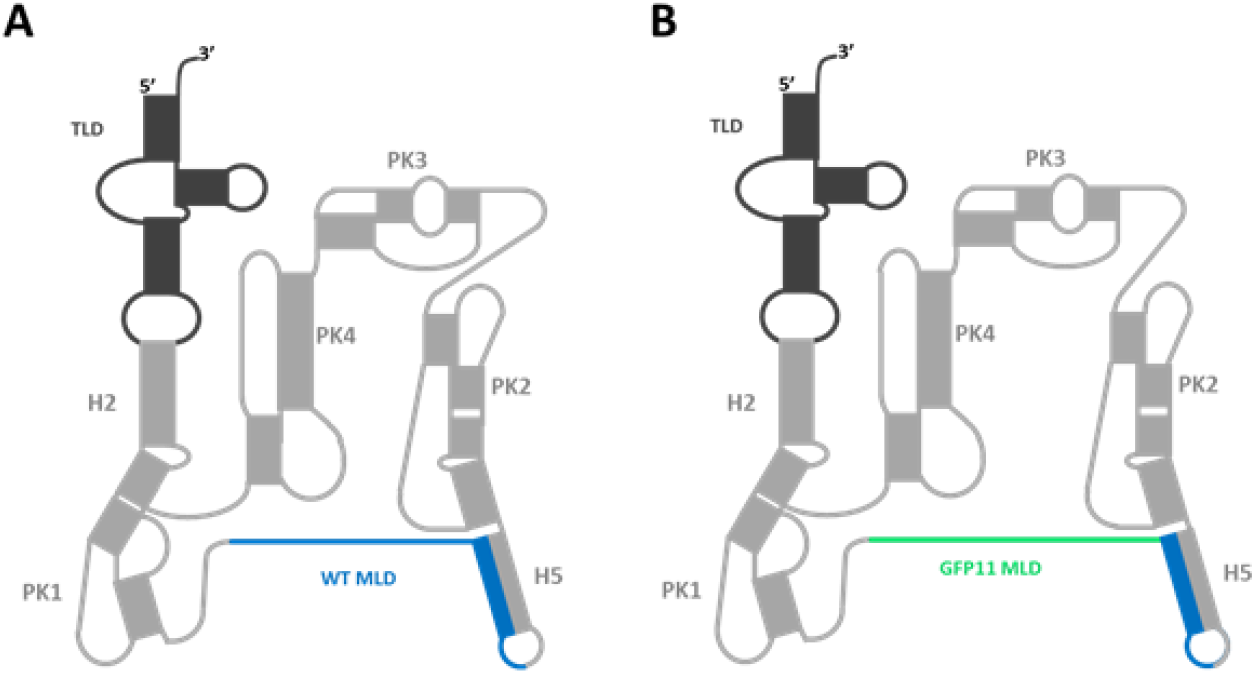
(A) Wild-type (WT) tmRNA. (B) Version 2 mutated tmRNA_GFP11_. An engineered MLD encoding the eleventh GFP domain is green, the WT 3’-end MLD is blue, and the WT H5 helix is gray.

**Supplementary Table 1:**
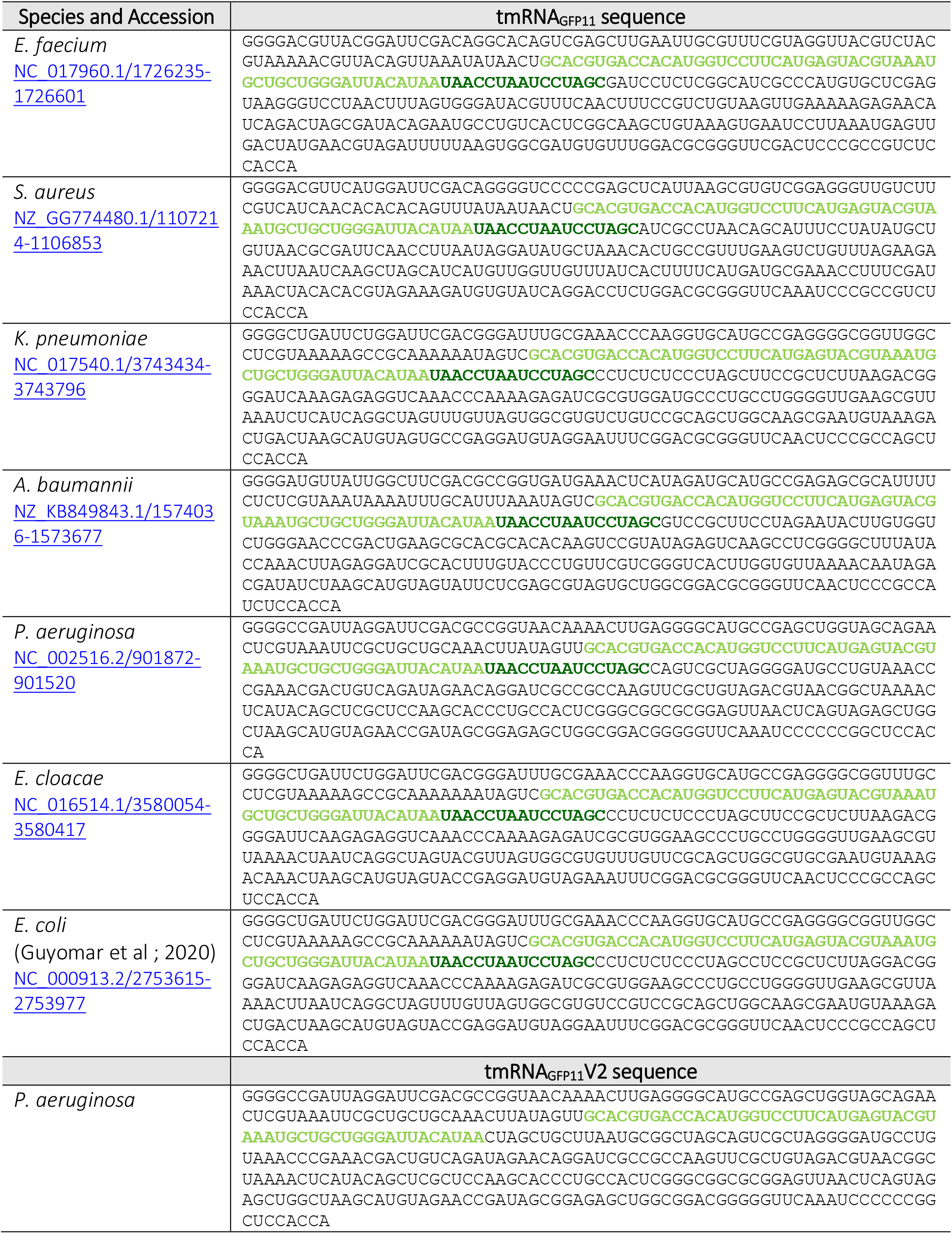
*E. coli* and ESKAPE tmRNA_GFP11_ sequences for *trans*-translation *in vitro.* The eleventh domain is light green and the compensatory mutations forming the helix H5 are darker green.

**Supplementary Table 2:**
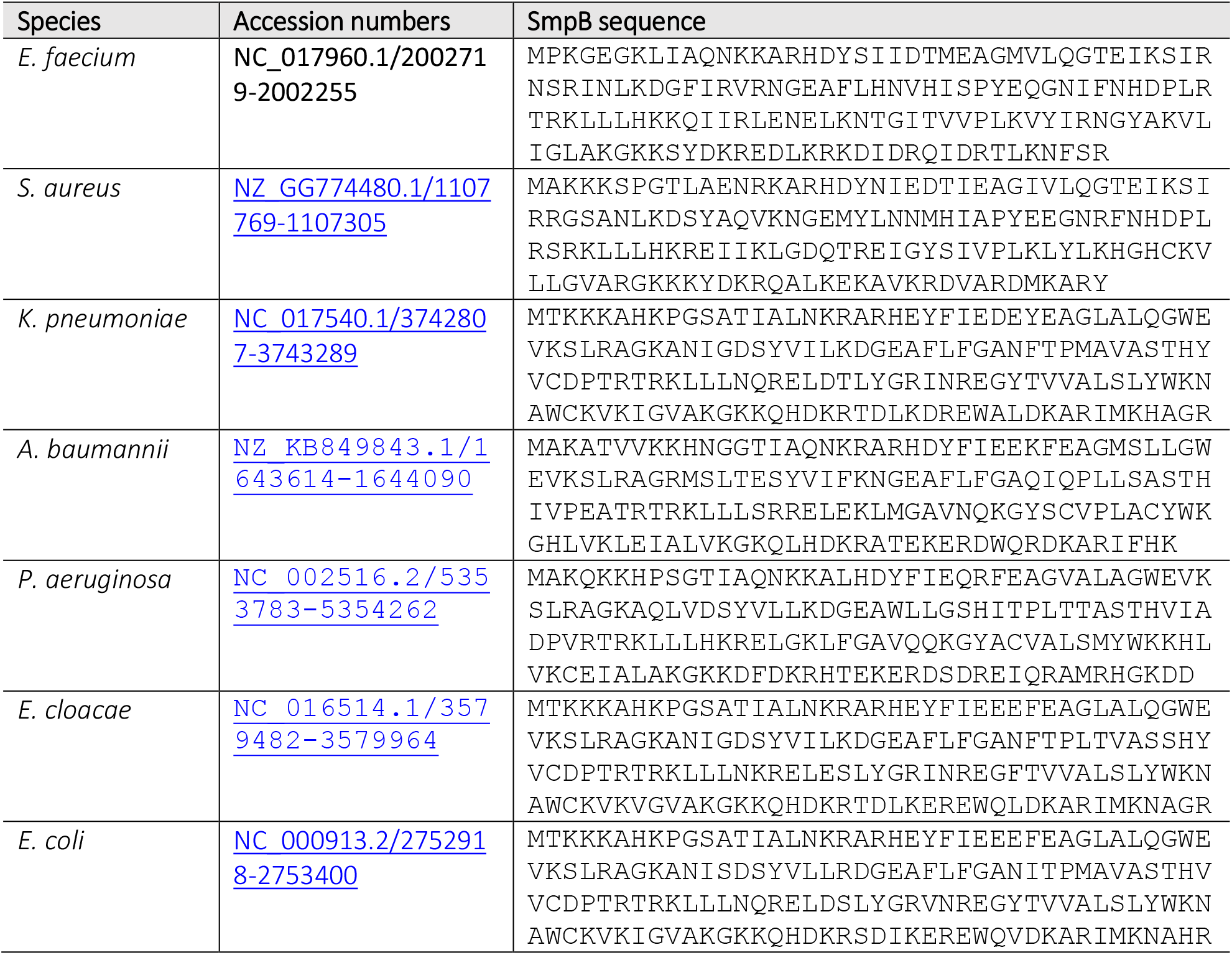
*E. coli* and ESKAPE SmpB amino acid sequences

**Supplementary Table 3:**
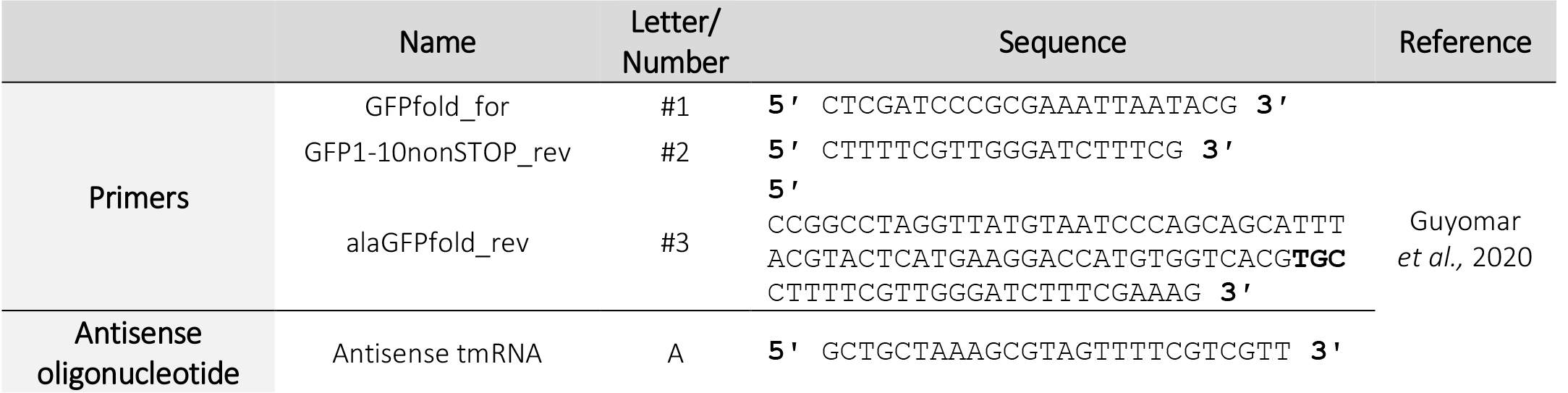
Primers and antisense oligonucleotide sequences.

**Supplementary Table 4:**
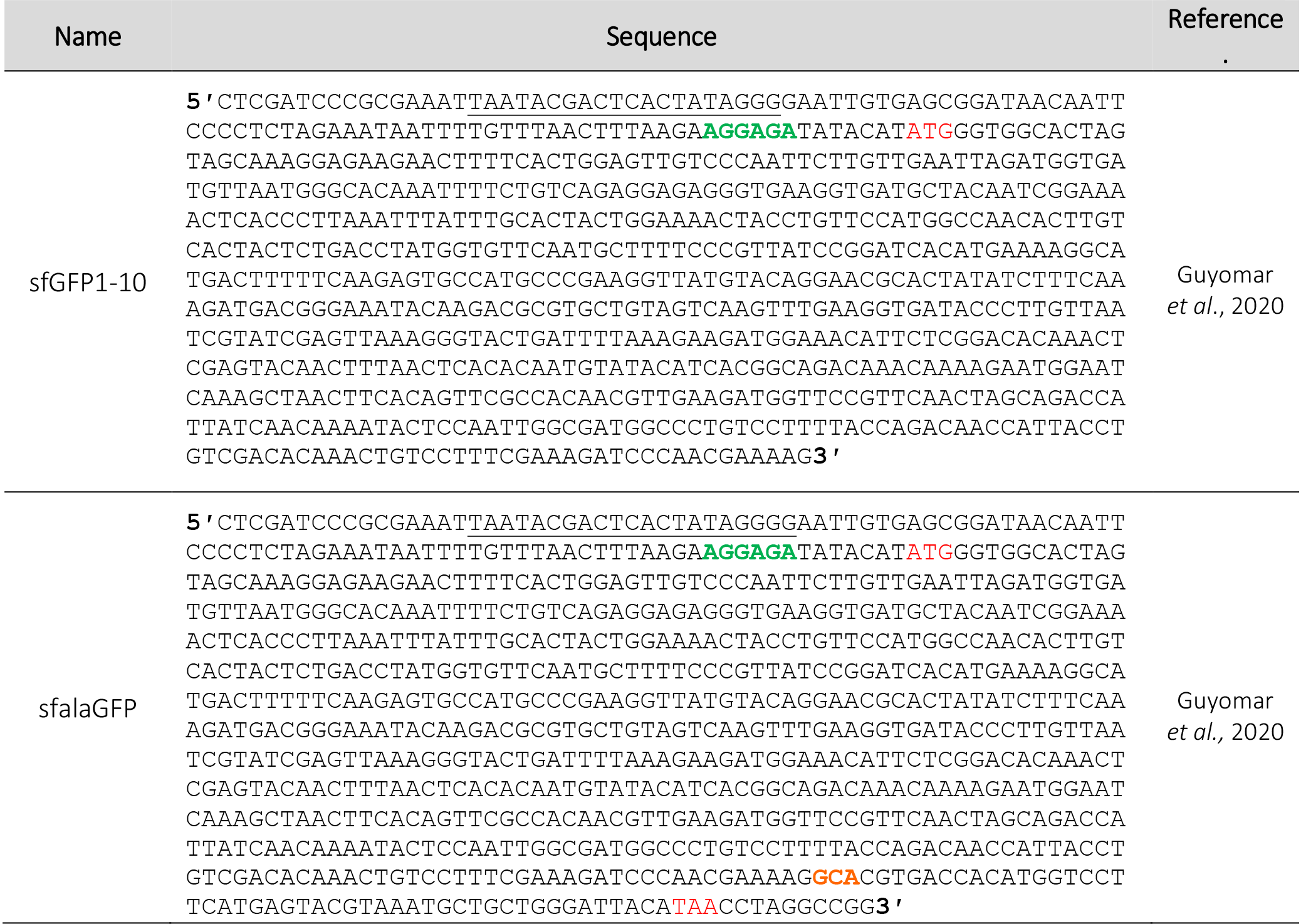
PCR product sequences for *in vitro trans*-translation. The T7 promoter is underlined, the RBS sequence is green, the start and stop codons are red, and the tmRNA alanine resume codon is orange.

**Supplementary Table 5:**
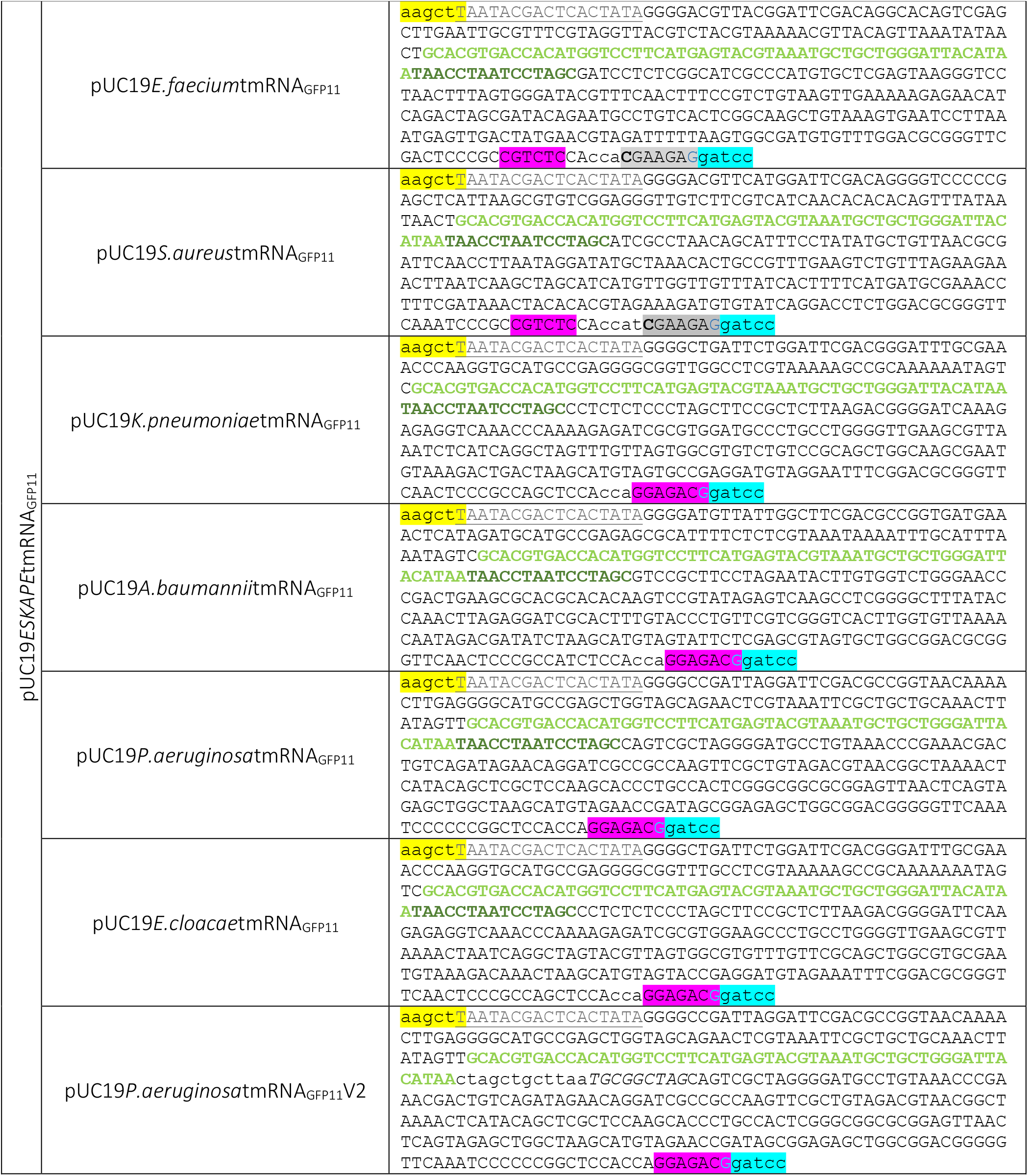
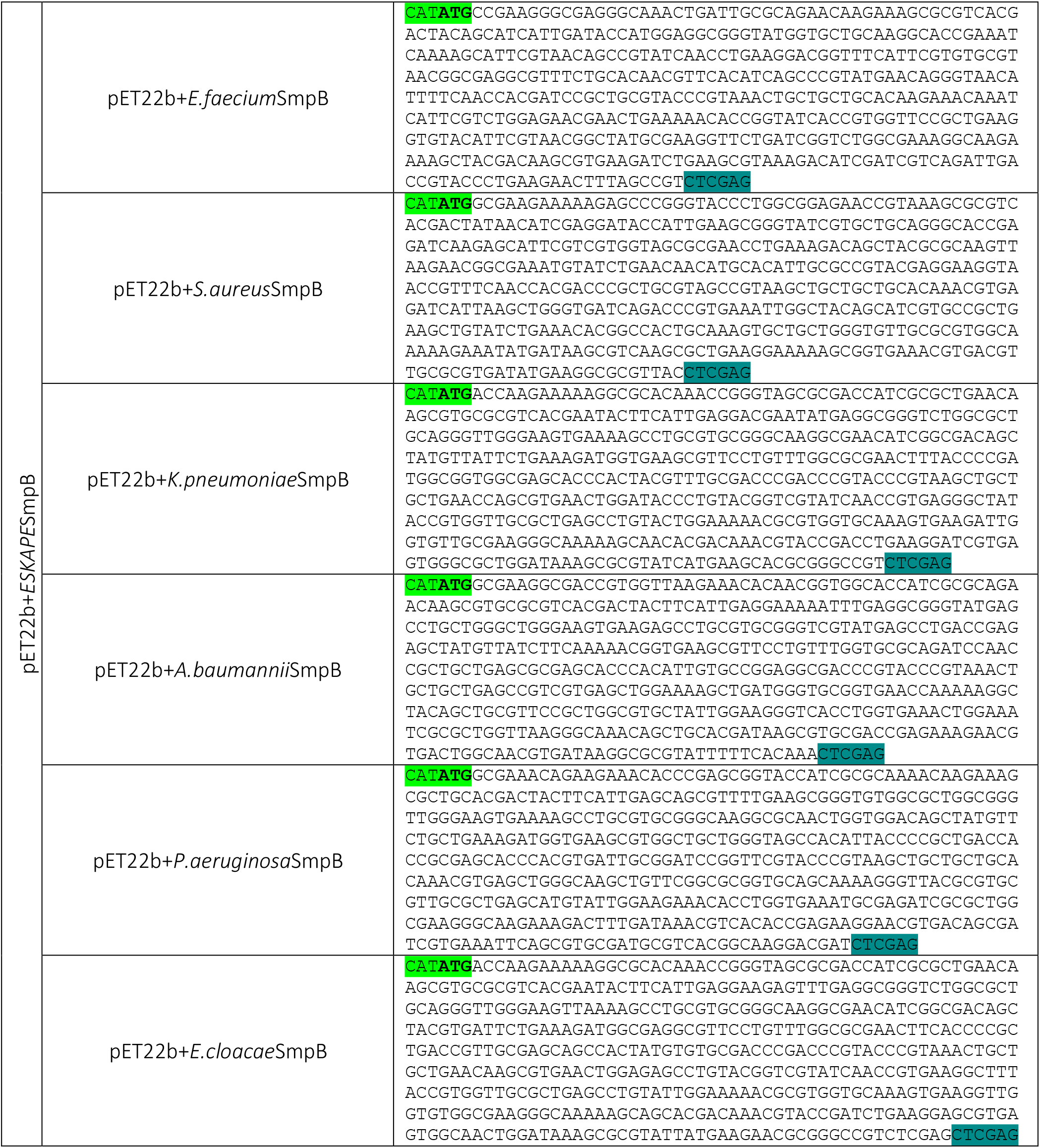
Plasmid list. These synthesized sequences were cloned by GenScript in pUC19 between the HindIII and BamHI restriction sites or in pET22b+ between the Ndel and Xhol restriction sites. The T7 promoter is grey and underlined, the eleventh domain is light green, and the mutations making up for the formation of the H5 helix are dark green. BsmB1 or EarI allow generation of the 3 ‘end of tmRNA. SmpB sequences are codon optimized for *E. coli.*

